# Th22 cells are a major contributor to the mycobacterial CD4+ T cell response and are depleted during HIV infection

**DOI:** 10.1101/732263

**Authors:** Rubina Bunjun, Fidilia M.A. Omondi, Mohau S. Makatsa, Tracey L. Müller, Caryn S.L. Prentice, Robert J. Wilkinson, Catherine Riou, Wendy A. Burgers

**Author notes:** **Correspondence**: Wendy Burgers, Institute of Infectious Disease and Molecular Medicine, Faculty of Health Sciences, University of Cape Town, Observatory 7925, South Africa.

## Abstract

HIV-1 infection substantially increases the risk of developing tuberculosis (TB). Some mechanisms, such as defects in the Th1 response to *Mycobacterium tuberculosis* (*M.tb*) in HIV-infected individuals have been widely reported. However, Th1-independent mechanisms also contribute to protection against TB. To identify a broader spectrum of defects in TB immunity during HIV infection, we examined IL-17 and IL-22 production in response to mycobacterial antigens in individuals with latent TB infection (LTBI) and HIV co-infection. Upon stimulating with mycobacterial antigens, we observed a distinct CD4+ T helper lineage producing IL-22 in the absence of IL-17 and IFN-γ. Th22 cells were present at high frequencies in response to mycobacterial antigens in blood and contributed up to 50% to the CD4+ T cell response to mycobacteria, comparable in magnitude to the IFN-γ Th1 response (median 0.91% and 0.55%, respectively). Phenotypic characterization of Th22 cells revealed that their memory differentiation was similar to *M.tb*-specific Th1 cells (*i.e*. predominantly early-differentiated CD45RO+CD27+ phenotype). Moreover, CCR6 and CXCR3 expression profiles of Th22 cells were similar to Th17 cells, while their CCR4 and CCR10 expression patterns displayed an intermediate phenotype between Th1 and Th17 cells. Strikingly, mycobacterial IL-22 responses were three-fold lower in HIV-infected individuals compared to uninfected individuals, and the magnitude of responses correlated inversely with HIV viral load. These data provide important insights into mycobacteria-specific T helper subsets and suggest a potential role for IL-22 in protection against TB during HIV infection. Further studies are needed to fully elucidate the role of IL-22 in protective TB immunity.

## INTRODUCTION

Tuberculosis (TB) is the leading cause of death from an infectious agent, claiming 1.6 million lives in 2017, with 10 million new TB cases that year (1). This considerable burden of disease, along with a host of challenges in diagnosing, treating and managing TB, emphasize its significance as a global health threat. Although TB is curable and successful treatment outcomes are typically >80%, cure is achieved less frequently with drug resistant TB (56%), and outcomes during HIV co-infection are worse (2). Importantly, cure does not lead to protection from re-infection or disease reactivation. HIV-infected persons are particularly vulnerable to developing TB, with an estimated increase in risk of 20-30 fold (3). The widespread introduction of ART has coincided with only a modest decline in TB in regions most affected by HIV (4), as TB risk still remains elevated in HIV-infected persons compared to HIV-uninfected persons, despite immune reconstitution (5).

The development of an effective TB vaccine is hampered by a lack of understanding of correlates of immune protection (6), particularly the functional and phenotypic characteristics of effector T cells that mediate control of *Mycobacterium tuberculosis* (*M.tb*), and how this immune response might be balanced by immunoregulatory T cell populations to limit inflammation and avoid pathology. The recent demonstration of the first candidate TB vaccine capable of protecting adults from pulmonary TB with an efficacy of 54% (7), provides the field with an opportunity to define correlates of vaccine protection, and has the potential to uncover unique insights into immunological control of TB.

TB and HIV co-infection presents us with a further prospect to improve our understanding of the mechanisms of immune control of *M.tb*, by identifying how HIV renders the immune response to *M.tb* defective, leading to increased risk of TB disease. CD4+ T cells and specifically the Th1/IFN-γ response to *M.tb* are critical for protective immunity to TB (8). Most studies of HIV-TB co-infection focus on Th1 immunity, and have demonstrated depletion or dysfunction of *M.tb*-specific Th1 responses in both blood (9–12) and the airways (13–15) during HIV infection.

However, there is evidence of a role for IFN-γ-independent mechanisms in immune control of TB (16) that may also contribute to, or synergize with, Th1 responses to TB. Recently, we characterized the profile of Th subsets specific for *M.tb* using lineage-defining transcription factors, revealing the broad spectrum of Th subsets involved in mycobacterial immunity, demonstrating that the inflammatory environment associated with HIV infection skewed these profiles (17). Th17 cells form part of this spectrum of *M.tb*-specific Th responses, and are believed to play an important role in immune protection from TB (18). Suppression of Th17-related genes was recently shown to be associated with progression to TB disease in *M.tb*-infected adolescents (19). In line with this, *M.tb*-specific IL-17-producing CD4+ T cells were significantly depleted in HIV-infected individuals from a TB-endemic area, compared to HIV-uninfected individuals (20).

Whilst IL-17 responses in *M.tb* immunity have been relatively well-studied (21–26), IL-22 responses have been overlooked in part due to their classification as a Th17 cytokine from studies in mice (27). In humans, however, IL-22 is produced by a distinct subset of CD4+ T cells (28–30), termed “Th22 cells”. IL-22 is a member of the IL-10 family of cytokines, and functions mainly to protect tissues from inflammation and infection, through stimulating proliferation and repair, and the production of antimicrobial peptides (31). Until recently, IL-22 was thought to be dispensable for control of *M.tb*, since deficiency or neutralization of IL-22 in mice had no effect control of *M.tb* using lab strains H37Rv and Erdman (32–35). However, the recent observation that IL-22 deficient mice infected with a clinical strain of *M.tb* (HN878) had an impaired ability to control *M.tb*, leading to increased bacterial burden and greater dissemination of infection (36), has triggered renewed interest in IL-22 and its role in TB control.

Given the paucity of data on *M.tb*-specific IL-22 CD4+ responses, and the knowledge that HIV infection results in the preferential targeting and depletion of Th22 cells (37), we sought to characterize HIV-induced defects in adaptive immunity to *M.tb*, with a focus on Th22 cells. Our findings highlight the large contribution IL-22 makes to the human CD4+ T cell response to TB (equivalent in magnitude to the IFN-γ response), with *M.tb*-specific Th22 cells being entirely distinct from Th1 and Th17 cells. Moreover, we show for the first time that *M.tb*-specific Th22 cells are depleted during HIV co-infection to a similar extent as Th1 responses. These findings emphasize the potential importance of this understudied CD4+ Th subset in TB immunity, and suggest that the loss of *M.tb*-specific Th22 cells may contribute to the increased risk of TB during HIV infection.

## MATERIALS AND METHODS

### Study Participants

Volunteers were recruited from Cape Town, South Africa, and fell within the following groups: ART naive HIV-seropositive persons with CD4 counts >400 cells/mm^3^ (n=25; median age 31; 96% female) and HIV-seronegative persons (n=25; median age 23; 60% female). HIV RNA levels were determined using an Abbott m2000 RealTime HIV-1 assay and blood CD4 counts by the Flow-CARE™ PLG CD4 test. All volunteers were TB sensitized based on a positive IFN-γ release assay (IGRA; Quantiferon, Cellestis), and active TB was excluded, based on symptoms and radiological evidence.

Healthy donors were recruited from the University of Cape Town, South Africa. Participants were >18 years of age, weighed >55 kg, did not have any chronic disease, did not use immunosuppressive medication and were not pregnant or lactating. These studies were approved by the Research Ethics Committee of the University of Cape Town (158/2010, 279/2012). All participants provided written, informed consent.

### Whole blood stimulation assays

Venous blood was collected and processed within 4 hours. Whole blood stimulation was performed as previously described (38) with the following antigens: Bacillus Calmette-Guerin (BCG; MOI of 4; SSI), Purified Protein Derivative (PPD) of *M. tuberculosis* (20μg/ml; Statens Serum Institute), ESAT-6 and CFP-10 peptide pools (4μg/ml), *M. tuberculosis* whole cell lysate (10μg/ml; BEI Resources) or PMA and Ionomycin (0.01μg/ml and 1μg/ml, respectively, Sigma), in the presence of anti-CD28 and anti-CD49d (1μg /ml each). Unstimulated cells were incubated with co-stimulatory antibodies only. Brefeldin A (BFA, 10μg/ml; Sigma) was added 7 hours after the onset of stimulation, and five hours after BFA addition, cells were either stained immediately, or red blood cells were lysed, the cell pellet stained with a violet viability dye, ViViD (Molecular Probes), fixed with FACS Lyse (BD Biosciences) and cryopreserved in 10% DMSO in FCS.

### Antibody Staining and Flow Cytometry

Cryopreserved or freshly stimulated whole blood was stained as previously described (15). For intracellular markers, cells were permeabilized with Perm/Wash buffer (BD Biosciences) and then stained intracellularly. Cells were stained with the following antibodies: CD3 APC-H7 (SK7; BD Biosciences), CD4 PE-Cy5.5 (S3.5; Invitrogen), CD4 ECD (T4; Beckman Coulter), CD8 QDot705 (3B5; Invitrogen), CD45RO ECD (UCLH1; Beckman Coulter), CD27 PE-Cy5 (1A4CD27; Beckman Coulter), CXCR3 PE-Cy7 (1C6/CXCR3; BD Biosciences), CCR6 BV605 (11A9; BD Biosciences), CCR4 BV510 (L291H4; Biolegend), CCR10 PE (1B5; BD Biosciences), KLRG1 PE-vio770 (REA261; Miltenyi Biotec), CD26 FITC (M-A261; BD Biosciences), IFN-γ Alexa700 (B27; BD Biosciences), IL-17 Alexa488 (N49-653; BD Biosciences), IL-17 FITC (BL-168; Biolegend), IL-22 PE (22URTI; e-Bioscience) or IL-22 eFluor450 (22URTI; e-Bioscience). Cells were acquired on a BD Fortessa using FACSDiva software and data analysed using FlowJo (TreeStar) and Pestle and Spice (39). A positive cytokine response was defined as twice background and a net response >0.025%, and all data are reported after background subtraction. A minimum of 30 cytokine-positive events was required for memory or chemokine receptor phenotyping.

### Statistical Analysis

Statistical analyses were performed using Prism 7 (GraphPad). Non-parametric tests (Mann-Whitney U test, Wilcoxon matched pairs test and Spearman Rank test) were used for all comparisons. Kruskal-Wallis with Dunn’s post-test was used for multiple comparisons. A p value of <0.05 was considered significant.

## RESULTS

### IL-22 responses are a major component of the CD4+ mycobacterial response

We examined CD4+ T cell cytokine profiles in response to a range of mycobacterial antigens in 25 healthy, HIV-uninfected persons sensitized by *M. tuberculosis* (*M.tb* IGRA+; **Table 1**). **Figure 1A** shows representative flow cytometry plots of IFN-γ, IL-22 and IL-17 CD4+ responses to *M. bovis* BCG, *M.tb* PPD and a pool of ESAT-6 and CFP-10 peptides from *M.tb*. As expected, CD4+ T cell IFN-γ responses to BCG were detected in all donors (median 0.55%, IQR: 0.28-1.46%; **Figure 1B**). Remarkably, IL-22 accounted for the greatest proportion of the CD4+ response to BCG (median 0.91%, IQR: 0.52-1.24%). In fact, the frequency of IL-22+ cells was greater than IFN-γ in 75% of participants. IL-17 CD4+ responses to BCG were significantly lower (median 0.11%, IQR: 0.06-1.66%) than both IFN-γ (p=0.007) and IL-22 (p=0.0008). Stimulation with *M.tb* PPD led to the detection of a similar IFN-γ response as BCG (median 0.74%), with comparatively lower frequencies of PPD-specific IL-22+ CD4+ T cells (median 0.21%; p=0.02) and IL-17+ cells (median 0%; p<0.0001) (**Figure 1B**). The ESAT-6/CFP-10 response was dominated by IFN-γ (median 0.07%), with low to undetectable IL-17 and IL-22 responses (medians of 0%). Taken together, these data demonstrate that different mycobacterial antigen preparations result in detection of different CD4+ T cell cytokine profiles. Of note, IL-22 made a substantial contribution to the anti-mycobacterial CD4+ response, with responses equivalent to or greater than the IFN-γ response to BCG.

**Table 1:**
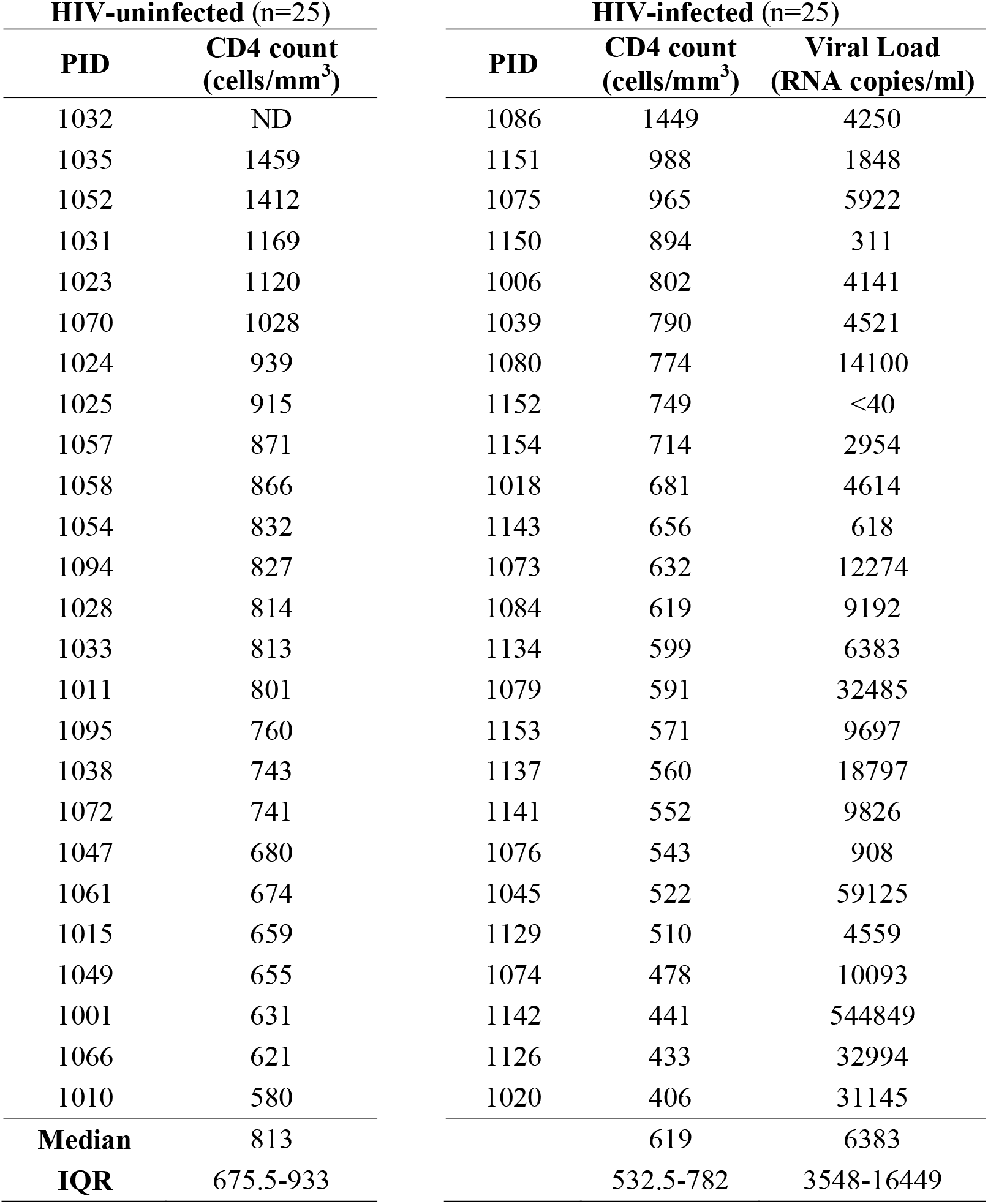
Characteristics of study participants

**Figure 1:**
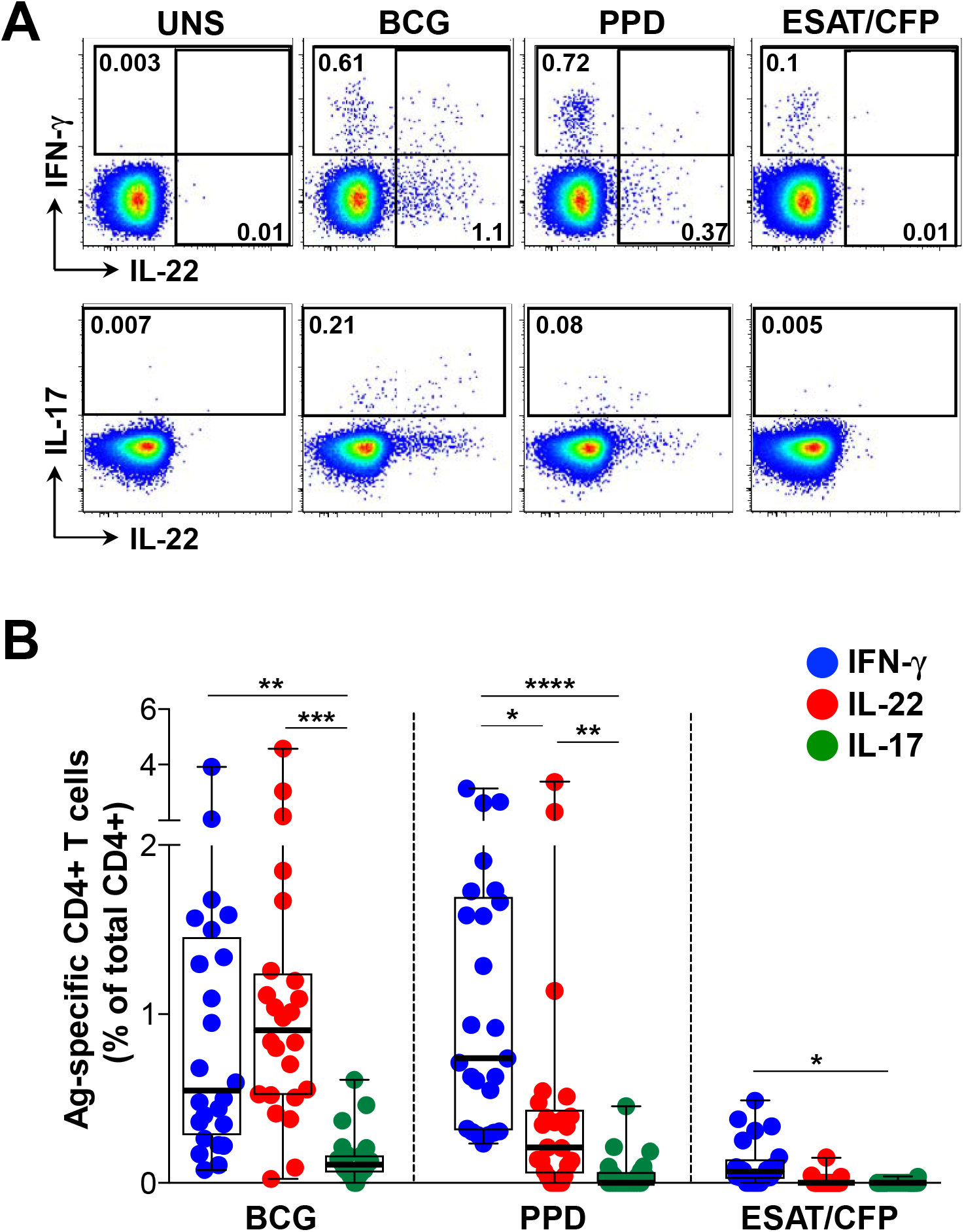
CD4+ T cell cytokine responses to mycobacterial antigens in latent TB infection. **(A)** Representative flow cytometry plots of the production of IFN-γ, IL-22 and IL-17 from CD4+ T cells after stimulation with *M. bovis* BCG, *M.tb* PPD and ESAT-6/CFP-10 peptides, in one study participant. UNS corresponds to the unstimulated control. The frequency of cytokine-producing cells is shown as a percentage of the total CD4+ T cell population, after gating on live, CD3+ lymphocytes. **(B)** Individual IFN-γ (blue), IL-22 (red) or IL-17 (green) responses to BCG, PPD or ESAT-6/CFP-10 (n=25). The frequency of cytokine-producing cells is shown as a percentage of the total CD4+ T cell population, after gating on live, CD3+ lymphocytes. Each dot represents one individual. Data are shown as box and whisker (interquartile range) plots and horizontal bars represent the median. Statistical comparisons were performed using a Kruskal-Wallis and Dunn’s multiple comparison test. *p≤0.05, **p≤0.01, ***p≤0.001, ****p≤0.0001

### Most CD4+ T cells producing IL-22 do not make IFN-γ and IL-17

We next focused on the high magnitude IL-22 response detected to BCG, to further characterize IL-22 CD4+ responses and their relationship with IFN-γ and IL-17. There was a highly significant positive correlation between IFN-γ and IL-22 responses to BCG (p<0.0001, r=0.830; **Figure 2A**). The frequency of IL-17+ CD4+ T cells also correlated with both IFN-γ and IL-22 production (p=0.039, r=0.424 and p=0.005, r=0.559, respectively; data not shown). Given these associations between IFN-γ, IL-22 and IL-17, we examined the co-expression patterns of the cytokines following BCG stimulation (**Figure 2B**). The majority of BCG-responding CD4+ T cells produced only IL-22 (median 47%; IQR: 36.6-59.6), whilst CD4+ cells secreting IFN-γ-alone made up a median of 37% (IQR: 27.1-47.4). There was minimal co-expression of IL-22 with both IL-17 (median 0.5%) and with IFN-γ (median 6.4%). When examining all CD4+ T cells producing IL-22, a median of 78% produced IL-22 alone (IQR: 71.1-89.2%), while 14% and 1.5% co-expressed IFN-γ or IL-17, respectively (data not shown). We also investigated IL-22 production in combination with other cytokines and found low or negligible co-expression with TNF-α, IL-2 and IL-21 (medians 0.3%, 0.5% and 3%, respectively, data not shown). Our data reveal that the large proportion of BCG-specific IL-22 was produced predominantly by CD4+ T cells secreting IL-22 in the absence of either IL-17 or IFN-γ, consistent with being a distinct ‘Th22’ lineage (28–30).

**Figure 2:**
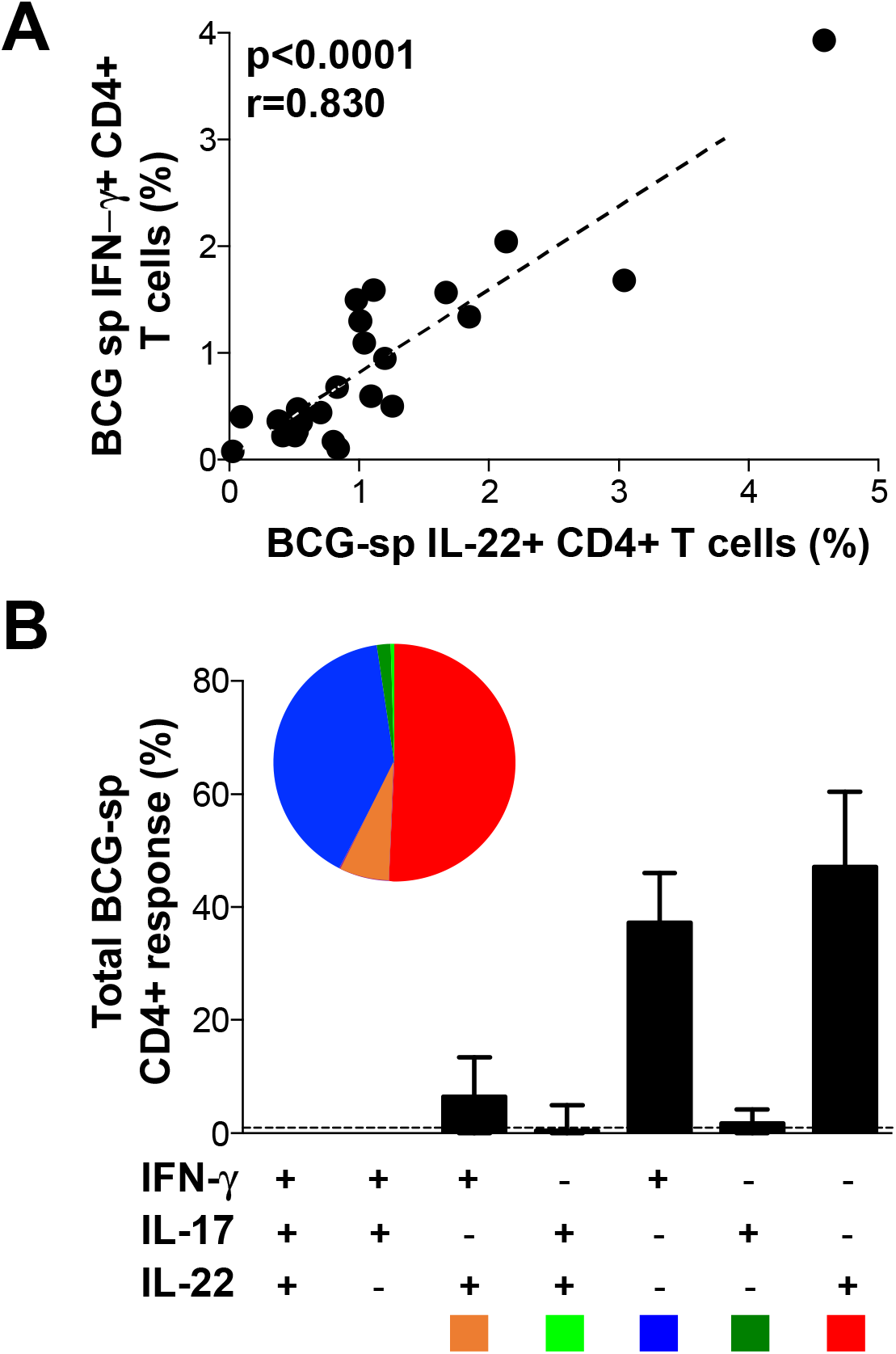
The relationship between IL-22, and other cytokines produced in response to BCG. **(A)** The relationship between the frequency of CD4+ T cells producing IFN-γ and IL-22 in response to BCG (n=24). Each dot represents an individual. Statistical analyses were performed using a non-parametric Spearman rank correlation. **(B)** Populations of CD4+ T cells producing different combinations of IFN-γ, IL-22 and IL-17 in response to BCG. The pie charts indicate the proportion of cytokine combinations that makes up the BCG response. Each slice of the pie represents a specific subset of cells, defined by a combination of cytokines shown by the color at the bottom of the graphs. Data are shown as box and whisker (interquartile range) plots and horizontal bars represent the median.

### Phenotypic characteristics of mycobacteria-specific IL-22-producing CD4+ T cells

In order to characterize the Th22 subset in more detail, we determined the memory differentiation profile of mycobacteria-specific Th22 cells (*i.e*. those producing IL-22 alone) compared to cells producing only IFN-γ or IL-17. **Figure 3A** shows representative flow cytometry plots of CD45RO and CD27 expression on total CD4+ cells with overlays of BCG-specific cytokine-producing CD4+ T cells (IFN-γ, IL-22 or IL-17 alone). The memory profile of BCG-specific CD4+ T cells was comparable, regardless of their cytokine secretion profile, with approximately 79% having an early differentiated phenotype (ED: CD45RO+CD27+, comprising central and transitional memory cells). Of the remaining cells, a median of ~17% were late differentiated (LD: CD45RO+CD27-, comprising effector memory and intermediate cells), with few terminally differentiated (TD: CD45RO-CD27-; ~0.3%) or naïve-like (CD45RO-CD27+; ~2 %) cells (**Figure 3B**). Thus, CD4+ T cells producing IFN-γ, IL-22 or IL-17 shared a similar memory differentiation phenotype.

**Figure 3:**
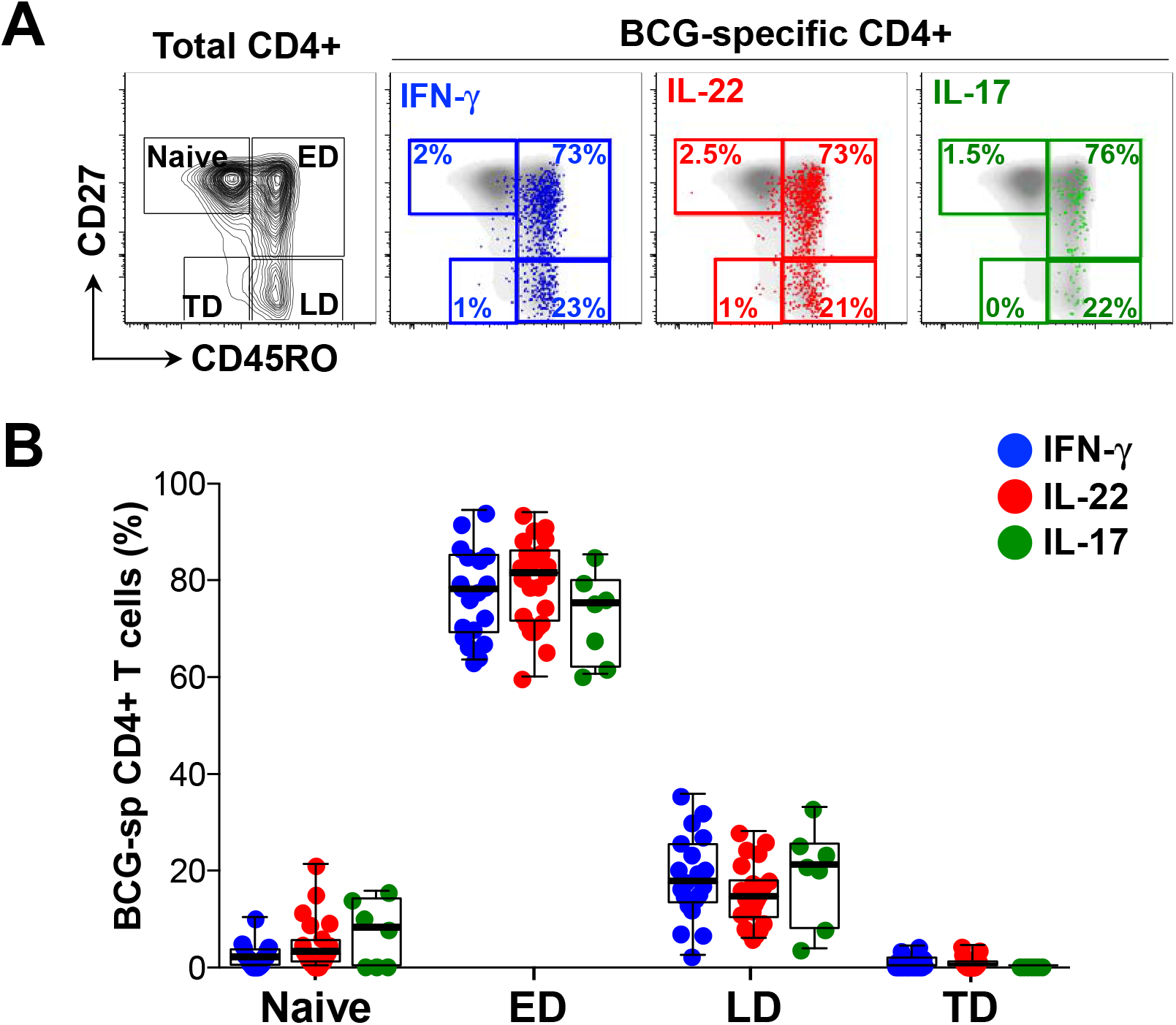
Memory profiles of CD4+ T cells producing IFN-γ, IL-22 or IL-17 in response to BCG. **(A)** Representative flow cytometry plots of total CD4+ memory subset distribution in one individual based on CD45RO and CD27 staining. Naïve: CD45RO-CD27+, early differentiated (ED: CD45RO+CD27+), late differentiated (LD: CD45RO+CD27-) and terminally differentiated (TD: CD45RO-CD27-). The overlays indicate the antigen specific CD4+ T cells producing IFN-γ (blue), IL-22 (red) or IL-17 (green). The frequencies of each subset are indicated. **(B)** The memory distribution of cells producing IFN-γ (blue), IL-22 (red) or IL-17 (green) in response to BCG (n=20, 25 and 7, respectively). Only individuals with a positive cytokine response and more than 30 cytokine events were included in the phenotyping. Each dot represents one individual. Data are shown as box and whisker (interquartile range) plots and horizontal bars represent the median. Statistical comparisons were performed using a Kruskal-Wallis and Dunn’s multiple comparison test.

To further characterize the phenotype of the different cytokine-producing subsets, we examined chemokine receptor expression profiles on CD4+ cells producing IFN-γ, IL-22 or IL-17. For these studies, we stimulated whole blood with *M.tb* whole cell lysate. To ensure that we were measuring similar cytokine responses, we compared IFN-γ, IL-22 and IL-17 induced by each antigen and found highly comparable frequencies of CD4+ T cell responses in the same donors (**Supplemental Figure S1**). **Figure 4A** shows representative flow cytometry plots of *M.tb*-specific CD4+ T cell production of IFN-γ, IL-22 and IL-17 overlaid onto chemokine receptor expression profiles (CXCR3, CCR6, CCR4 and CCR10). Whilst a majority of IFN-γ-producing cells expressed CXCR3 (median 61.5%), a sizable fraction also expressed CCR6 (median 54.5%), with a low proportion expressing CCR4 (median 4.1%) and negligible CCR10 (median 0.5%; **Figure 4B**). In contrast, CD4+ cells producing IL-22 were almost all CCR6 positive (median 94.7%), and compared to cells producing IFN-γ, significantly fewer expressed CXCR3 (median 27.7%), and significantly more expressed CCR4 and CCR10 (median 23.3% and 2.7%, respectively). Th17 cells (IL-17+) shared comparable expression profiles for CCR6 and CXCR3 (medians 96.1% and 17.2%, respectively), but a higher proportion expressed CCR4 (median 50.8%) and CCR10 (median 5.8%) compared to Th22 cells. Of note, cells co-producing IFN-γ and IL-22 had a similarly high expression of CCR6 as Th22 and Th17 cells, but were otherwise intermediate between IFN-γ+ and Th22 for the remaining chemokine receptors (**Supplemental Figure S2A**). These findings demonstrate distinct patterns of chemokine receptor expression on different cytokine-producing subsets. These data are consistent with previous descriptions (28, 30), but also highlight the substantial overlap in chemokine receptor expression between T helper subsets producing distinct cytokines.

**Figure 4:**
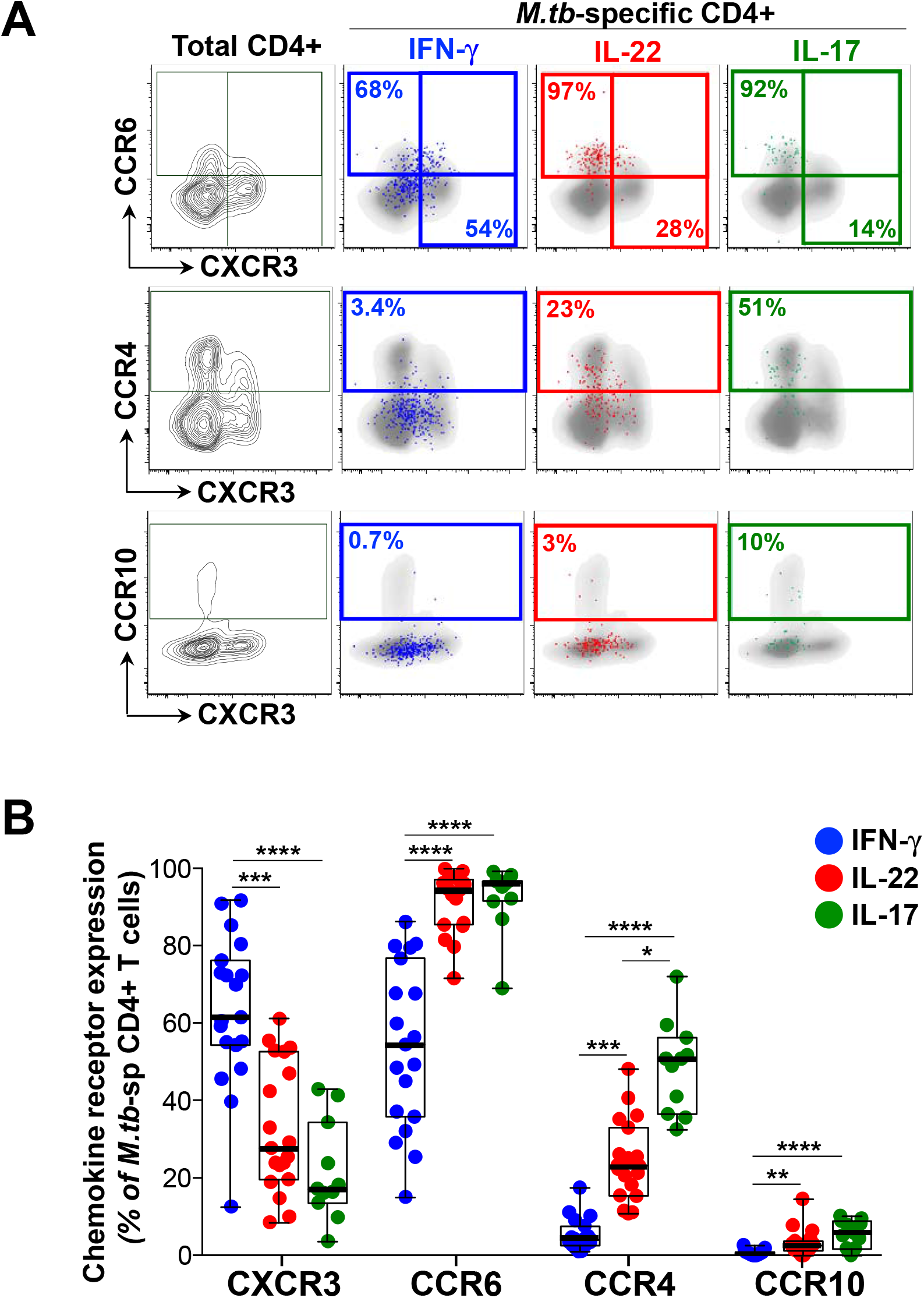
Chemokine receptor expression of CD4+ T cells producing IFN-γ, IL-22 or IL-17 in response to *M.tb* whole cell lysate. **(A)** Representative flow cytometry plots of the expression of CCR6, CCR4, CXCR3 and CCR10 on total CD4+ T cells in one individual. The overlays indicate the antigen specific CD4+ T cells producing IFN-γ (blue), IL-22 (red) or IL-17 (green). The frequencies of each subset are indicated. **(B)** The chemokine receptor distribution of cells producing IFN-γ (blue), IL-22 (red) or IL-17 (green) in response to *M.tb* lysate (n=19, 19 and 11, respectively). Only individuals with a positive cytokine response and more than 30 cytokine events were included in the phenotyping. Each dot represents one individual. Data are shown as box and whisker (interquartile range) plots and horizontal bars represent the median. Statistical comparisons were performed using a Kruskal-Wallis and Dunn’s multiple comparison test. *p≤0.05, **p≤0.01, ***p≤0.001, ****p≤0.0001

We also investigated the homing potential of *M.tb*-specific CD4+ Th subsets using KLRG1 and CD26. Killer cell lectin-like receptor G1 (KLRG1)-expressing cells appear to be retained within lung blood vasculature, while KLRG1^−^ cells migrate to the lung parenchyma (40). Dipeptidyl peptidase IV (CD26) is involved in enzymatic chemokine modification that enhances T cell migration (41, 42). Expression of these markers was significantly different between Th subsets (**Supplementary Figure S2B**). Th22 cells were characterized by a near absence of KLRG1 expression compared to Th1 and Th17 cells. In contrast, 50% of Th22 cells expressed CD26, compared to a median of 34% of Th1 cells and 11% of Th17 cells (**Supplemental Figure S2B**). These data suggest that *M.tb*-specific Th22 are endowed with a distinct homing potential compare to Th1 and Th17 cells.

### The effect of HIV infection on the Th22 response to mycobacteria

Th1 responses to *M.tb* are impaired or reduced during HIV infection (3). However, little is known about the effect of HIV co-infection on the Th22 response to *M.tb*. Hence, we examined IFN-γ, IL-22 and IL-17 responses to BCG and PPD in 25 HIV-infected individuals with a median CD4 count of 619 cells/mm^3^ (IQR: 532.5-782) and a median plasma viral load of 6.38 ×10^3^ copies/ml (IQR: 3.55-16.45 ×10^3^; **Table 1**). Consistent with previous reports, the frequency of *M.tb*-specific CD4+ T cells producing IFN-γ was significantly lower in HIV-infected participants compared to uninfected participants in response to BCG (p=0.0004, medians 0.12% and 0.55%, respectively; **Figure 5A**). Notably, the IL-22 response to BCG was also lower in HIV-infected individuals, to a similar degree as the IFN-γ response (p=0.0005; medians 0.28% and 0.91%, respectively). Additionally, IL-17 responses were also significantly lower in HIV-infected individuals in response to BCG compared to the HIV-uninfected group (p<0.0001, medians 0% and 0.11%, respectively). After adjusting for CD4 count, these differences became even more evident (**Figure 5B**), despite the relatively well-preserved CD4+ T cell numbers in our HIV-infected cohort. HIV-infected participants had 8-fold (p<0.0001) and 3-fold (p=0.0003) fewer CD4+ T cells producing IFN-γ or IL-22, respectively, compared to uninfected participants. There were also fewer cells producing IL-17 in HIV-infected individuals (median 0; p<0.0001). Similar results were obtained for IFN-γ and IL-22 in response to PPD (**Supplemental Figure S3A and B**). Overall, HIV-infected participants had lower *M.tb*-specific IFN-γ, IL-22 and IL-17 responses. Whilst the decrease in *M.tb*-specific IFN-γ and IL-17 responses during HIV infection has been reported, we report here a striking loss of *M.tb*-specific CD4+ T cells producing IL-22.

**Figure 5:**
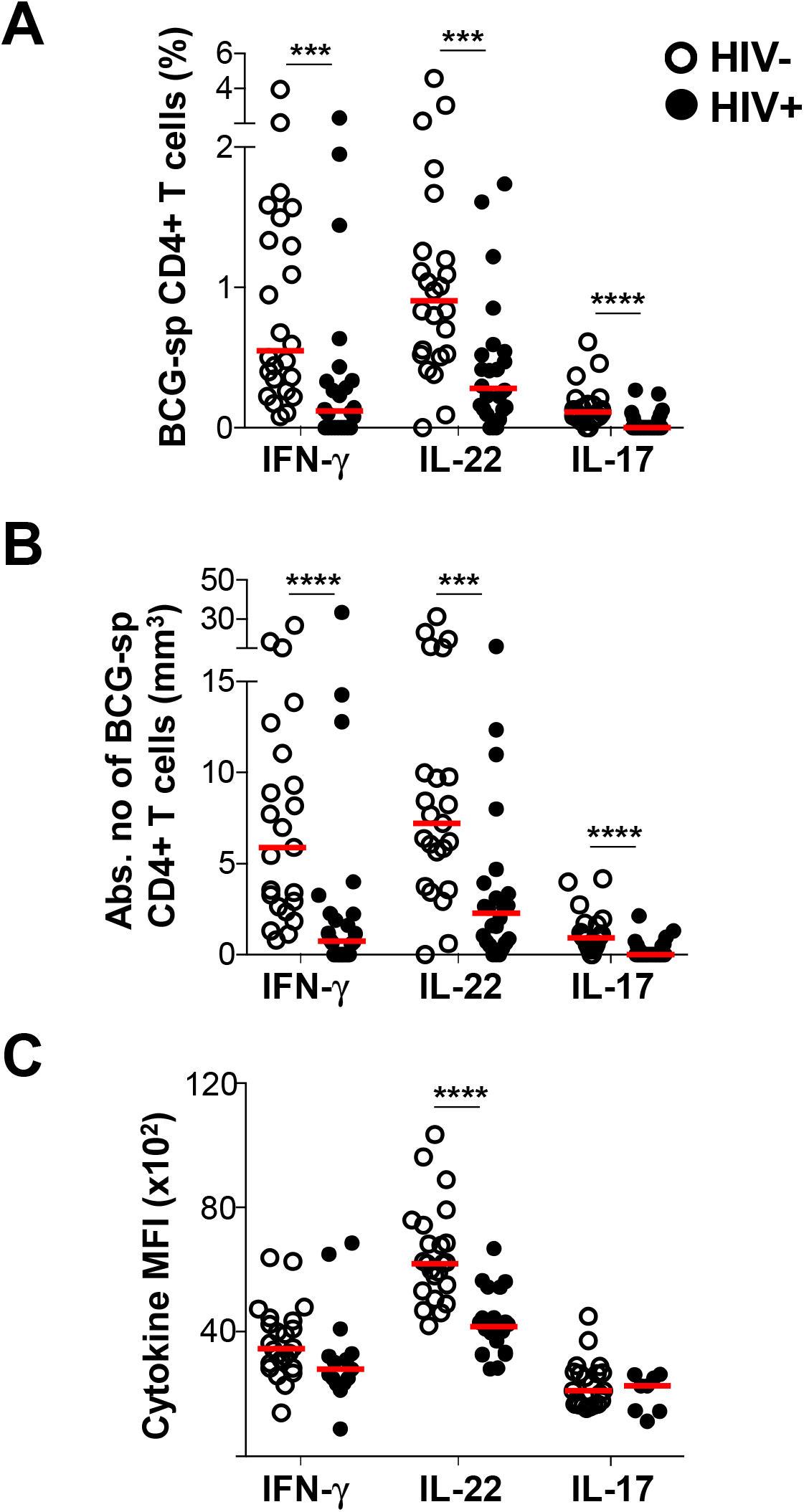
CD4+ T cell responses to BCG in HIV-infected and uninfected individuals. **(A)** The individual IFN-γ, IL-22 or IL-17 responses in HIV uninfected or infected individuals in response to BCG (n=24 in each group). **(B)** The cytokine frequency adjusted for CD4 count in HIV-infected and HIV-uninfected individuals in response to BCG. **(C)** The median fluorescent intensity (MFI) of IFN-γ in response to BCG (n=24 and n=15 for HIV-uninfected and infected, respectively) The MFI of IL-22 in response to BCG (n=23 and n=20 for HIV-uninfected and infected, respectively) The MFI of IL-17 in response to BCG (n=22 and n=8 for HIV-uninfected and infected, respectively). For each cytokine, MFI was only graphed for individuals with positive cytokine responses. HIV-uninfected participants are shown with open circles and HIV-uninfected individuals with closed circles. Each dot represents one individual. Data are shown as box and whisker (interquartile range) plots and horizontal bars represent the median. Statistical comparisons were performed using a non-parametric Mann Whitney test. *p≤0.05, **p≤0.01, ***p≤0.001, ****p≤0.0001

To further investigate the impact of HIV on BCG-specific Th22 responses, we measured the amount of IL-22 produced per cell, using median fluorescent intensity (MFI). The MFI of IL-22 was significantly lower in HIV-infected individuals compared to uninfected individuals (p<0.0001; medians 4169 and 6215, respectively; **Figure 5C**), whereas no differences in the MFI of IFN-γ and IL-17 was observed. This suggests that HIV may have a unique effect on Th22 cells in response to BCG. However, we found no differences in the MFI of any cytokines produced in response to PPD (**Supplemental Figure S3C**).

To investigate whether the lower cytokine responses to mycobacterial antigens in HIV-infected individuals related to clinical parameters, the association between IFN-γ, IL-22 and IL-17 responses and CD4 count or plasma viral load was examined. We observed a significant positive correlation between both the IFN-γ and IL-22 response to BCG and CD4 count (p=0.04, r=0.43; and p=0.004, r=0.57, respectively; **Figure 6A**). Likewise, in response to PPD, IFN-γ (p=0.03, r=0.45) and IL-22 (p=0.004, r=0.57) correlated directly with CD4 count (data not shown). This suggests that the decrease in these responses could be a consequence of overall CD4+ T cell depletion, despite the relatively narrow CD4 count range and modest CD4 decreases in our study (84% of participants had CD4 counts >500 cells/mm^3^). No association between the frequency of IL-17 and CD4 count was observed for either BCG (**Figure 6A**, bottom panel) or PPD (data not shown). Finally, there were no significant associations between plasma viral load and IFN-γ or Th17 responses to BCG; **Figure 6B**) or any cytokine in response to PPD (data not shown). However, the frequency of Th22 cells responding to BCG was significantly inversely correlated with plasma viral load (p=0.006, r=−0.54, **Figure 6B**, middle panel).

**Figure 6:**
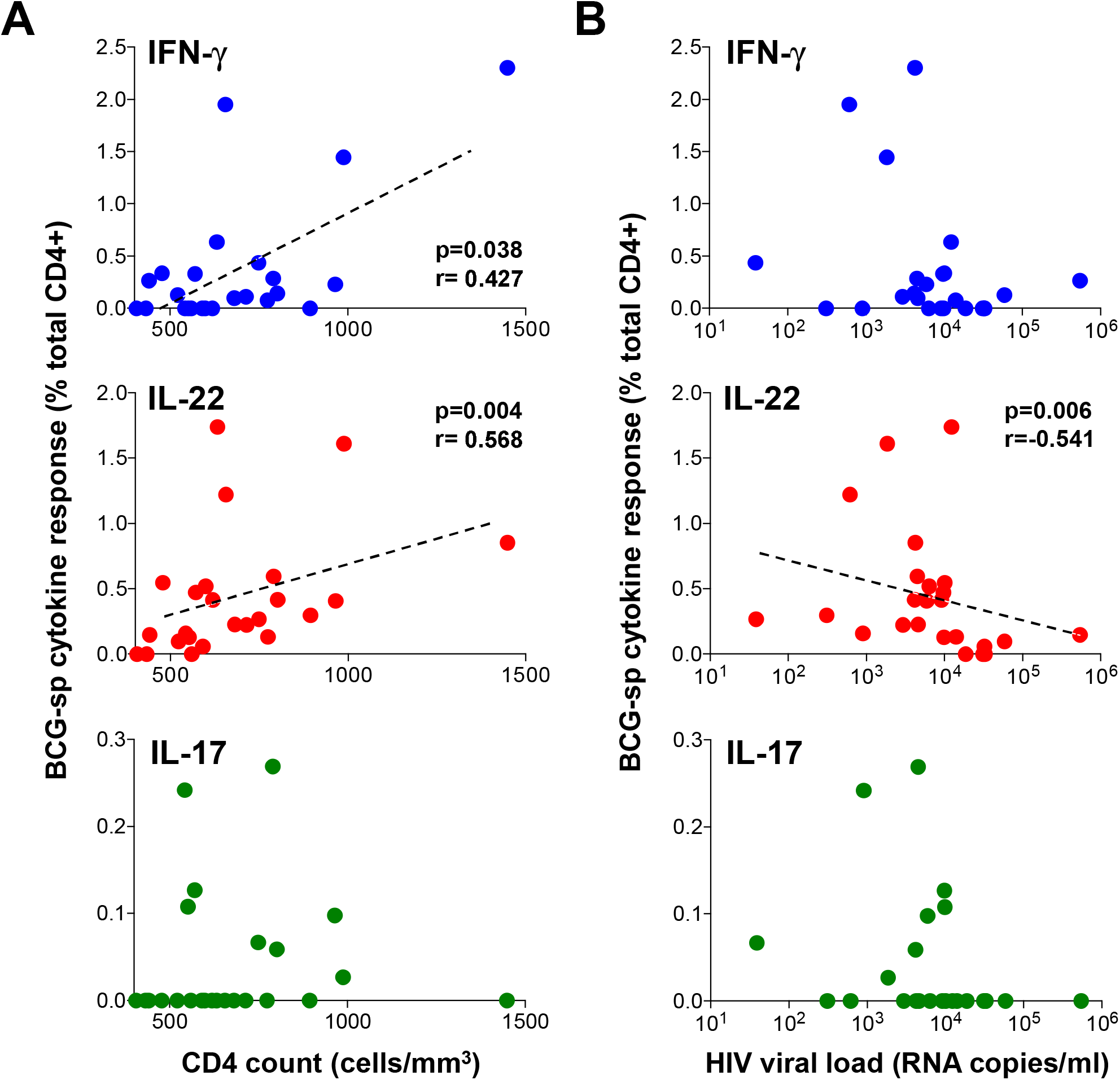
The relationship between clinical parameters and IFN-γ, IL-22 or IL-17 CD4+ T cell responses to BCG in HIV-infected individuals. The association between IFN- (blue), IL-22 (red) or IL-17 (green) responses to BCG and **(A)** CD4 count or **(B)** viral load. Each dot represents an individual (n=24). The dotted line indicates linear regression for statistically significant correlations, highlighted in bold. Statistical analyses were performed using a non-parametric Spearman rank correlation.

Overall, we demonstrate the detrimental effect of HIV infection on CD4+ T helper subsets in response to mycobacteria. In particular, the Th22 subset exhibited both a decrease in the magnitude of the response to mycobacteria, and a defect in IL-22 production on a per cell basis. Furthermore, unlike the other cytokine-producing subsets examined (Th1 and Th17), the frequency of Th22 cells correlated inversely with HIV viral load, suggesting a direct relationship between HIV infection and the loss of Th22 cells specific for mycobacteria.

## DISCUSSION

Th1/IFN-γ responses are needed for an effective response to TB (8), however a range of immune mechanisms beyond Th1 immunity may also contribute to protection from TB (6). Since HIV-infected individuals are considerably more susceptible to TB disease (3), key components required for effective immune control of *M.tb* are likely to be defective in these individuals, and we sought to identify these. In addition to IFN-γ/Th1 immunity, this study examined IL-17 and IL-22 responses to mycobacteria in *M.tb*-sensitized, HIV-infected and uninfected individuals. Consistent with previous studies, we identified distinct populations of CD4+ T cells expressing IFN-γ, IL-17 or IL-22 in response to mycobacterial antigens (43, 44). The IL-22 response was unexpectedly abundant, contributing up to 50% of the mycobacterial response measured using these three cytokines, and the source was a distinct subset of CD4+ T cells producing IL-22 alone. Importantly, IL-22 response was impaired in HIV-infected individuals in both magnitude and function, suggesting that depletion of this subset may contribute to TB risk.

IL-22 has classically been characterized as a Th17-related cytokine, since in mice it is co-secreted with IL-17 and has overlapping functions with IL-17 (27). However, IL-22 is a member of the IL-10 family (45), and in humans IL-22 is not co-expressed with IL-17 (28–30). Consequently, ‘Th22’ cells were proposed as a novel CD4+ T helper cell lineage in humans, with shared but distinct features and functions compared to Th17 cells. To date, the role of IL-17 in *M.tb* immunity has been well-studied (21–24, 26). Here, we found that IL-17 responses made only a modest contribution to the total mycobacterial response in *M.tb*-exposed individuals, consistent with previous reports (20, 43). In contrast, we detected ample mycobacteria-specific IL-22 production from CD4+ T cells in the absence of IL-17 (and IFN-γ), consistent with a distinct Th22 subset and in agreement with earlier observations in LTBI and TB disease (43, 46). Phenotypic profiling demonstrated that whilst their memory differentiation phenotype was similar to that of Th1 and Th17 cells, the bulk of Th22 cells expressed CCR6, with expression frequencies of CXCR3, CCR4 and CCR10 intermediate between Th1 and Th17 cells, somewhat consistent with published reports (28, 30). *M.tb*-specific Th22 cells were also characterized by higher CD26 and absent KLRG1 expression compared to both Th1 and Th17 cells. Altogether, these characteristics emphasize the shared and unique features of mycobacteria-specific Th22 cells relative to Th1 and Th17 cells, which may relate to distinct homing capabilities.

The previously unappreciated, sizeable contribution Th22 cells make to the mycobacterial response prompts the question of whether Th22 responses play a role in protective immunity against *M.tb*. Previous studies demonstrated that deficiency or neutralization of IL-22 in mice did not affect control of laboratory strains of *M*.tb (H37Rv and Erdman) (32–35). However, renewed interest in IL-22 has been garnered since the observation that IL-22 deficient mice infected with a hypervirulent clinical strain of *M.tb* (HN878) have an impaired ability to control the infection, resulting in both increased bacterial burden and greater dissemination of infection (36). Additional evidence from a range of models suggest that IL-22 may indeed participate in TB immunity. IL-22 has been found at sites of TB disease; soluble IL-22 and IL-22 transcripts were elevated in the airways, lung tissue, granuloma, and in pleural and pericardial effusions during TB disease (43, 46–50). Along with IFN-γ, IL-22 was one of the strongest genes upregulated in bovine TB (51), and gene expression signatures revealed that IFN-γ and IL-22 were the dominant correlates of protection from bovine TB in blood in BCG-vaccinated cattle (52). Human genetic studies demonstrated the association between increased susceptibility to TB and a single nucleotide polymorphism in the IL-22 promoter that decreased IL-22 expression (50).

If IL-22 is involved in TB immunity, how might it mediate a protective function? IL-22 functions as a key regulator of tissue-specific antimicrobial immunity (31). The receptor for IL-22 is a heterodimer consisting of the IL-10R2 and the IL-22R, and expression is primarily restricted to non-hematopoietic cells, particularly epithelial cells in the skin, digestive tract and respiratory tract (31). IL-22 has been shown to be essential for mediating protective immunity to a range of extracellular and intracellular bacteria, such as *Klebsiella* and *Chlamydia* in the lung and *Citrobacter* in the intestine (53–56). Neutralization of IL-22 led to bacterial dissemination, exacerbated pathology, and lower Th1 and Th17 responses in the lung (55). The protective role at barrier sites appears to be mediated by three distinct functions, namely; maintenance of barrier integrity by promotion of epithelial homeostasis, stimulating epithelial proliferation and preventing apoptosis, as well as enhancing mucin production and tight junction formation; inducing antimicrobial peptides such as β-defensins; and regulating chemokine secretion from epithelial cells to co-ordinate recruitment of immune cells, such as neutrophils, to inflamed tissue (27, 29, 55, 57). Indeed, Treerat and colleagues demonstrated that the TB-protective role of IL-22 resulted from the secretion of S100 and Reg3γ from epithelial cells, and induction of CCL2 that mediated macrophage recruitment to the infected lung (36). It is worth noting that several studies have independently documented IL-22R expression on *M.tb*-infected monocyte-derived macrophages (MDMs), as well as macrophages in TB granulomas in humans and non-human primates (36, 58, 59). Consistent with these findings, IL-22 from CD4+ T cells and innate cells, as well as recombinant IL-22, reduced mycobacterial replication in MDMs by improving phagolysosome fusion (36, 58–60). These data suggest that a direct effector function for IL-22 in limiting mycobacterial growth cannot be ruled out.

HIV-infected individuals remain one of the most vulnerable populations at risk of TB (3). The early depletion of *M.tb*-specific Th1 responses, considered fundamental to TB immunity, has been reported during HIV infection (9, 10). Here, we investigated the relative effect of HIV on Th22 and Th17 responses to mycobacteria compared to Th1 responses. An important and novel finding from our study was that the mycobacteria-specific Th22 response was depleted during HIV infection, to a similar extent as Th1/IFN-γ responses. Several studies have described a global and preferential loss of Th22 and Th17 cells during HIV/SIV infection, leading to mucosal gut damage and systemic immune activation, driving HIV disease progression (37, 61–64). The CCR6+CD4+ T cell subset (within which all Th22 and Th17 cells reside) is more permissive to HIV infection and replication, and is enriched for HIV DNA (65–67). Elevated expression of HIV co-receptors CCR5 and CXCR4 has been reported on CCR6+CD4+ T cells, which could facilitate HIV entry (68). In addition, post-entry mechanisms appear to create a more permissive cellular environment for HIV replication in CCR6-expressing cells, demonstrated by specific transcriptional signatures favoring HIV replication (69–71). We report here that higher HIV plasma viral load correlates with lower frequencies of Th22 cells specific for mycobacteria, consistent with a mechanism of direct, preferential infection of Th22 cells by HIV. Overall, multiple mechanisms may contribute to the loss of Th22, Th17 and Th1 subsets specific for *M.tb* (72, 73), and their combined depletion may contribute to TB risk during HIV infection.

Our new findings add to a growing body of evidence in support of a role for IL-22 in protective immunity to TB. However, a number of questions remain unanswered. Does IL-22 contribute to protective immunity to TB, or only during infection with specific clinical strains, or during HIV infection, when multiple immunological defects manifest? Does IL-22 assume a direct effector or indirect regulatory role in immunity to TB, or both? Does the inflammatory context dictate whether IL-22 might be beneficial to the host or pathological (74)? Ultimately, will it be necessary to induce Th22 responses for a TB vaccine to be effective? Notwithstanding these gaps in our knowledge, our study highlights the substantial contribution that Th22 cells make to mycobacterial immunity, and the importance of further elucidating the role of IL-22 in the control of *M.tb* infection and disease.

## ACKNOWLEDGEMENTS

We thank the study participants for providing samples and for their time and commitment to the study, and to the clinical staff at the Ubuntu HIV-TB clinic. We thank Dr Shaun Barnabas and Rene Goliath for phlebotomy. We thank Mrs Kathryn Norman for administrative assistance. We are grateful to BEI Resources, NIAID, NIH, for the following reagent: *Mycobacterium tuberculosis*, Strain H37Rv, Whole Cell Lysate, NR-14822.

## AUTHOR CONTRIBUTIONS

Conceived and designed the experiments: WAB, CR and RJW. Performed the experiments: RB, FMAO, MSM, TLM and CSLP. Analyzed the data: RB, FMAO, SMM and WAB. Wrote the paper: RB and WAB. All authors approved the final manuscript.

---

**Supplemental Figure S1: Cytokine responses to BCG and *M.tb* whole cell lysate.** Comparison of the frequencies of CD4+ T cells producing IFN-γ, IL-17 and IL-22 in response to BCG (circles) and *M.tb* whole cell lysate (triangles) in healthy donors (n=8). The frequency of cytokine-producing cells is shown as a percentage of the total CD4+ T cell population, after gating on live, CD3+ lymphocytes. Statistical comparisons were performed using a non-parametric matched pairs Wilcoxon test.

**Supplemental Figure S2: Phenotypic profiles of Th subsets. (A)** The chemokine receptor distribution of cells co-producing IFN-γ and IL-22 (n=14) in response to *M.tb* lysate. Horizontal bars represent the median. **(B)** KLRG1 and CD26 expression on *M.tb*-specific Th1, Th22 and Th17 cells. Representative overlay plots showing KLRG1 and CD26 expression on total CD4+ T cells (grey), IFN-γ+ (blue), IL-22+ (red) and IL-17+ (green) cells in response to *M.tb* lysate (top panel). Expression of KLRG1 and CD26 shown as box and whisker (interquartile range) plots (bottom panel) and horizontal bars represent the median (n=19). Statistical comparisons were performed using a Kruskal-Wallis and Dunn’s multiple comparison test. Only individuals with a positive cytokine response and more than 30 cytokine events were included in the phenotyping. Each dot represents one individual.

**Supplemental Figure S3: CD4+ T cell responses to PPD in HIV-infected and uninfected individuals. (A)** The individual IFN-γ, IL-22 or IL-17 responses in HIV uninfected (n=25) or infected individuals (n=24) in response to PPD. **(B)** The cytokine frequency adjusted for CD4 count in HIV-infected and HIV-uninfected individuals in response to PPD. **(C)** The median fluorescent intensity (MFI) of IFN-γ in response to PPD (n=25 and n=24 for HIV-uninfected and infected, respectively). The MFI of IL-22 in response to PPD (n=21 and n=18 for HIV-uninfected and infected, respectively). The MFI of IL-17 in response to PPD (n=12 and n=12 for HIV-uninfected and infected, respectively). For each cytokine, MFI was only plotted for individuals with positive cytokine response. HIV-uninfected participants are shown with open circles and HIV-uninfected individuals with closed circles. Each dot represents one individual. Data are shown as box and whisker (interquartile range) plots and horizontal bars represent the median. Statistical comparisons were performed using a non-parametric Mann Whitney test.

