## Supplementary figures and images for "Th22 cells are a major contributor to the mycobacterial CD4+ T cell response and are depleted during HIV infection"

### Supplemental Figures 1-3

**FIGURE S1**
**Bunjun *et al.***

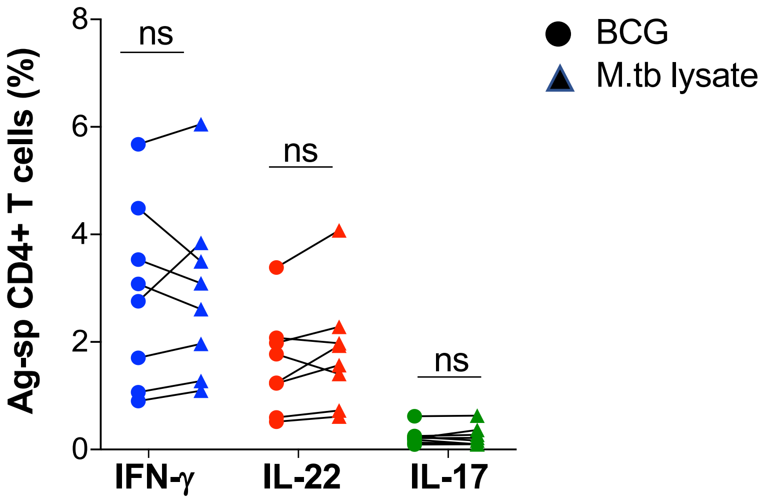

FIGURE S2
Bunjun *et al.*

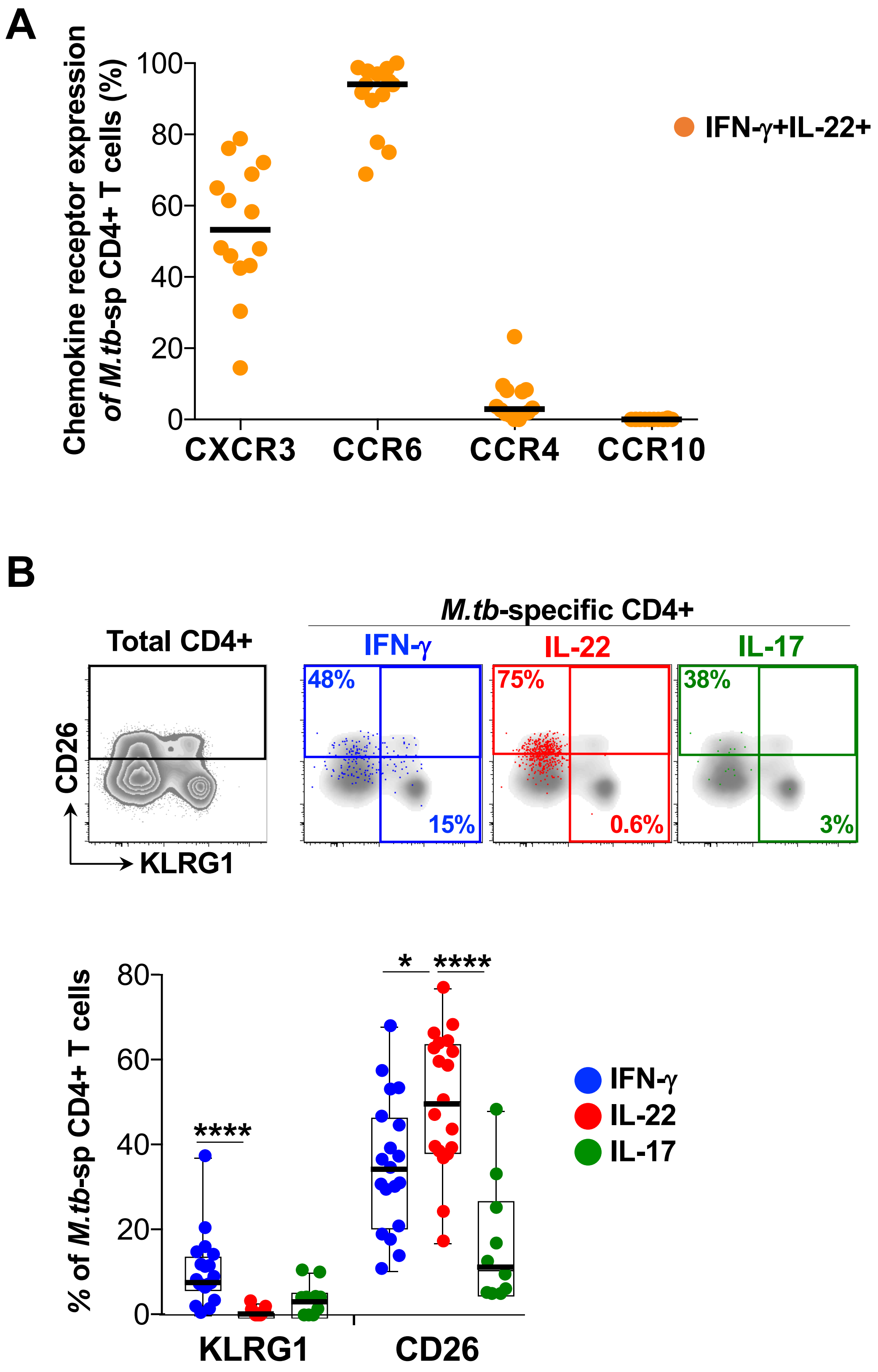

**FIGURE S3**
**Bunjun *et al.***

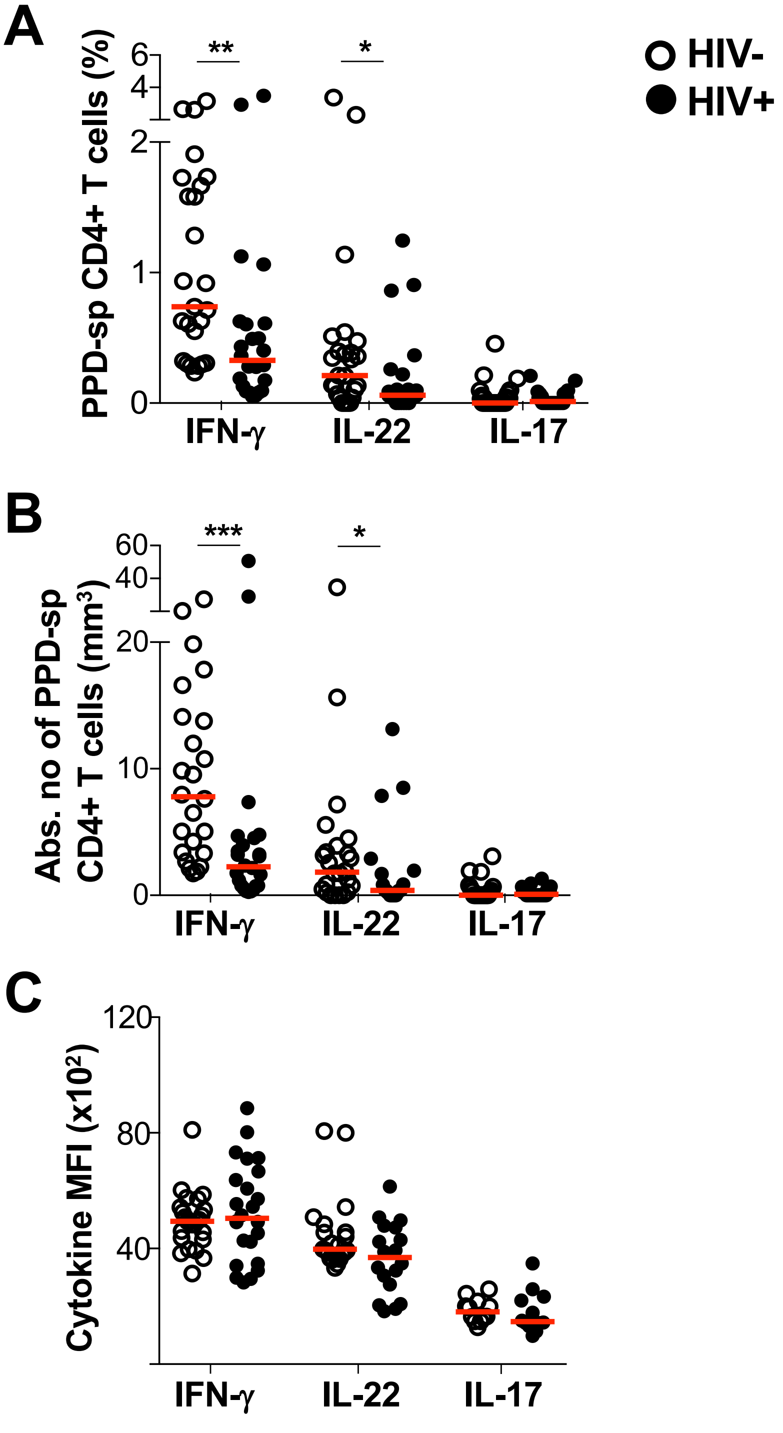
